# Disruption without diversity loss: how spreading thermotolerant *Pilidium lythri* infection perturbs plant-soil microbiome stability

**DOI:** 10.1101/2025.01.28.633866

**Authors:** Dominika Siegieda, Jacek Panek, S. Emilia Hannula, Magdalena Frąc

**Author notes:** correspondence: Dominika Siegieda.

## Abstract

While global warming accelerates phytopathogen spread, still little is known on how fungal pathogens disrupt fragile balance in plant-microbe interactions. Here, we demonstrate that the thermotolerant fungus *Pilidium lythri* combines broad chemical resistance with a remarkable ability to destabilize microbial communities in strawberry soil-plant systems. Using high-throughput sequencing (16S rRNA and ITS markers) across 72 samples, combined with mixed-effects modelling, source-tracking, microbiota assembly mechanisms investigation and network analysis we demonstrate that *P. lythri* disrupted bacterial community assembly processes, transitioning from deterministic to stochastic dynamics in the bulk soil, aligning with the Anna Karenina Principle. Despite stable dominance index (ENS) under infection bacterial and fungal communities, rhizosphere fungal diversity increased under infection. Network analysis of this niche revealed decreased fungal connectivity, with enhanced antagonistic interactions between taxa. We also noted that microbial migration patterns between niches under pathogen inoculation highlighted shifts in community interactions. Our work uncovers how emerging thermotolerant fungal phytopathogens destabilize plant holobiont without reducing diversity, important for managing emerging agricultural threats in a warming climate.

## Introduction

Fungal and bacterial microorganisms, together with their host plants, are interconnected through intricate, multi-level interactions that shape ecosystems and sustain plant health and productivity. Based on these relationships, the plant holobiont concept was formed - an integrated system where hosts and microbiota function as one. Plants produce root exudates to modulate microbial assembly of their niches and vicinity, while the microbiome, in turn, is able to influence and regulate plant health, improving its resilience against abiotic and biotic stressors (Berg, 2009; Baldrian, 2016; Trivedi *et al*., 2020; Gupta *et al*., 2021; Marco *et al*., 2022; Michl *et al*., 2023). These finely tuned interactions are widely studied and hold great potential for improving agricultural productivity in a sustainable manner (Trivedi *et al*., 2020; Vishwakarma *et al*., 2020; Frąc *et al*., 2022).

However, this delicate balance between microbiomes and plants is being increasingly threatened by ongoing global warming. The rise of annual global temperature and changed precipitation patterns has created the opportunity for a spread and establishment of invasive phytopathogens into new regions (Bebber *et al*., 2013; Helfer, 2014; Liu *et al*., 2019). Although substantial attention has been given to beneficial plant–microbe interactions, far less is known about how invasive or thermotolerant fungal pathogens perturb microbial community structure, niche interactions and assembly rules. This knowledge gap is particularly concerning for perennial crops like strawberries (*Fragaria x ananassa* Duch.), which depend heavily on stable rhizosphere and soil microbial communities to thrive and are particularly vulnerable to such disruptions.

Ecological theory offers a conceptual lens for understanding such perturbations. The *Anna Karenina Principle* (AKP) predicts that under biotic or abiotic stress, host-associated microbiomes become more variable and stochastic and unpredictible. Increasing evidence suggests that pathogen-induced disruption can generate divergent microbiome states rather than uniform shifts, but experimental demonstrations in soil–plant systems remain limited (Arnault *et al*., 2023).

*Pilidium lythri* is an emerging fungal phytopathogen with a versatile host range and limited genomic data. This ascomycete was originally reported in countries located in warmer climate zones, such as India (Phatak and Payak, 1965), Iran (Arzanlou *et al*., 2013; Sayami *et al*., 2013) and China (Zhang *et al*., 2008; Duan *et al*., 2010). Notably, its occurrence had been recently identified in several temperate climate countries, including Belgium (Debode *et al*., 2011), Croatia (Milicević *et al*., 2014), Spain (Aguín *et al*., 2018), France (Cardin *et al*., 2009; Caesar *et al*., 2012) and Poland, although in 2004 only morphologically, but in 2020 we confirmed its presence in symptomatic Polish strawberry fields with molecular methods (Gołębniak and Jarosz, 2004; Malarczyk *et al*., 2020), and with new isolates collected in present study. Primary targets of this fungal phytopathogen are strawberry plants and fruits (Lopes *et al*., 2010; Geng *et al*., 2012; Ayoubi *et al*., 2016), but it also affects other agriculturally significant plants, including *Olea europaea* L. (Arzanlou *et al*., 2013), *Cornus mas* (Torbati *et al*., 2019), *Prunus domestica* (Sayami *et al*., 2013), *Eucalyptus* spp. (Maússe-Sitoe *et al*., 2016), *Vitis vinifera* (Aguín *et al*., 2018), and various ornamental flowers (Zhang *et al*., 2008; Cardin *et al*., 2009; Caesar *et al*., 2012; Bruckart *et al*., 2013; Garfinkel and Chastagner, 2019). Notably concerning is that *P. lythri* is not host-specific. Strains isolated from olives were able to cause disease symptoms on strawberry fruits (Karimi *et al*., 2016), further increasing the severity of the threat posed by this pathogen as it spreads to new regions. Studies conducted in Brazil, Iran and Belgium showed incidence as high as 70% in high humidity and warm temperatures, highlighting its threat in warming climate (Lopes *et al*., 2010; Ayoubi *et al*., 2016; Karimi *et al*., 2016). Pathogenicity test revealed rapid progression of the disease, with symptoms appearing on up to 100% of fruits within days after inoculation (Debode *et al*., 2011; Fernández-Ortuño *et al*., 2014). The pathogen is able to infect both wounded and intact grape berries, further underscoring its high virulence (Aguín *et al*., 2018). Yet, no study has examined how *P. lythri* or other spreading, thermotolerant phytopathogens influence microbial processes in soil–plant interfaces or whether its effects produce AKP-like disruptions in microbial assembly.

Here, we hypothesised that infection by a thermotolerant fuganal pathogen destabilizes microbial community assembly across soil-plany niches, increasing stochasticity and altering microbial interaction patterns consistent with the Anna Karenina Principle. To test this, we (I) explored chemical astress tolerance of *P. lythri* across temperatures, (II) quantified pathogen-induced changes in bacterial and fungal communities in bulk soil, rhizosphere and roots using high-throughput sequencing, and (III) assessed assembly mechanisms, source-sink dynamics and network stability under infection. This approach contribute to a broader understanding how spreading pathogens influence plant-microbiome systems in the face of environmental change.

## Methods

### Fungal isolates and molecular identification

Pure fungal isolates were obtained by placing symptomatic strawberry leaves and fruits (surface-sterilized with detergent and 70% ethanol) on PDA medium and incubating them at 24°C in darkness for 7 days. After initial screening, fungi with macroscopic and microscopic characteristics matching the description of *P. lythri* (Palm, 1991; Marin-Felix *et al*., 2017; Karimi *et al*., 2019) were identified through amplification and Sanger sequencing of the Internal Transcribed Spacer (ITS) regions of fungal rDNA, with ITS1 and ITS4 primers (White *et al*., 1990), as previously described (Frąc *et al*., 2015). The obtained sequences were deposited in GenBank under accession numbers: PQ632564.1, PQ632565.1, PQ632566.1, PQ632567.1. These sequences were then used to construct phylogenetic tree with other sequences available in Genbank database from the *Pilidium* genus. Evolutionary analyses were conducted in MEGA X (Kumar *et al*., 2018). Initial tree for the heuristic search was obtained automatically by applying Neighbor-Join and BioNJ algorithms to a matrix of pairwise distances estimated using the Tamura-Nei model, and then selecting the topology with superior log likelihood value. The tree was drawn to scale, with branch lengths measured in the number of substitutions per site. This analysis involved 57 nucleotide sequences and there were a total of 372 positions in the final dataset. To fulfil Koch’s postulates, a spore suspension of 1.6 × 10^4^ spores/mL was prepared, and surface-sterilized strawberry leaves were briefly submerged in the solution. The leaves were then sealed in ziplock bags and incubated at 24°C in darkness for 7 days. Sterile deionized water was used as the untreated control.

### Metabolic profile arrays

Each cell of Phenotype Microarray 21-25 BIOLOG plates (Biolog, Hayward, CA, USA) was inoculated with the fungal mycelium as recommended by the manufacturer and described before (Panek *et al*., 2016). The plates are coated with 4 different concentrations of the chemical substances and precise concentrations of the substances used for microarrays are confidential and protected as proprietary information by the manufacturer, thus, for the purposes of this article, we refer to them as concentration levels 1, 2, 3, and 4 to maintain clarity. For each of 120 tested chemical substances (Supplementary Table 1), we incubated 3 *P. lythri* strains across four concentrations at two temperatures (24 and 30°C), resulting in 72 replicates per substance (3 strains x 3 replicates x 4 concentrations x 2 temperatures). The temperature of 24°C reflects the optimal *in vitro* growth conditions for *P. lythri*, while 30°C was selected to simulate elevated temperature scenarios associated with ongoing climate warming. This design allowed us to assess the thermal plasticity of the fungus’ metabolic response under environmental shifts. All plates were incubated for 96 hours in the darkness. Microplate reader (Biolog, Hayward, CA, USA) was used to determine the absorbance at 490 nm at the beginning of the experiment and every 24 hours which reflects the reduction of tetrazolium dye, portraying a metabolic activity of the analysed microorganism. Higher values indicate robust redox activity and potential tolerance against tested chemical. Obtained data was analysed statistically and visualised in RStudio, using tidyverse (Wickham *et al*., 2019) and lme4 (Bates *et al*., 2015) packages. First, values obtained at 0 h for each cell was divided from read values at 96 h to establish growth of metabolic activity, and the data was rescaled from 0 to 1. To establish which factors had an influence on fungal activity, we used generalized linear mixed-effects model (GLLM) with Gamma family, controlling for the plate. And to establish whether the substance inhibit fungus’ respiration rates, we controlled for plate, strain, temperature and concentration.

### Pot experiment and isolation of microbial DNA

Strawberry plants, obtained from the nursery, were grown in pots containing sieved soil from a certified organic strawberry plantation. After 20 days of initial growth of strawberry plants at 29°C and 70% humidity, half of the pots were inoculated with *P. lythri* G322/23 spores using 10 mL of a 2 · 10^4^ spores mL^-1^ solution per plant. Sterile, deionized water was used as a negative control. Samples were collected from two pots in triplicates (6 samples per niche per time point) for healthy and pathogen-inoculated samples in two timepoints: at the time of inoculation and again after 9 days of incubation. Bulk soil was transferred into sterile 1.5 mL Eppendorf tubes and frozen at −20°C. The rhizosphere was collected as described by Lundberg et al. (Lundberg *et al*., 2012). Roots, flushed free of rhizosphere soil, were also frozen at −20°C prior to DNA isolation which was conducted with the QIAQUBE instrument (Qiagen, Hilden, Germany) and GeneMATRIX Soil DNA Purification Kit (EURx, Gdańsk, Poland) after homogenization of the samples in the FastPrep-24 instrument (MP Biomedicals) at 4 m s^-1^ for 15 s. DNA isolates were then frozen in −20°C until library construction and sequencing.

### Metataxonomic sequencing, bioinformatics and statistical analysis

For the metataxonomic sequencing, microbial ITS2 (Schoch *et al*., 2012) and 16Sv3 regions (Klindworth *et al*., 2013) were amplified, indexed and sequenced on Illumina MiSeq instrument (2×300). Raw data was deposited at SRA database under accession numbers: PRJNA1213298 for 16S and PRJNA1213307 for ITS. Raw data was then passed through the custom QIIME2 (Bolyen *et al*., 2019) pipelines as reported before (Siegieda *et al*., 2023, 2024). In brief, after primer trimming with Cutadapt (Martin, 2011) and ITSxpress (Rivers *et al*., 2018), denoising, chimera removal, merging and Amplicon Sequence Variant (ASV) calling were performed. The taxonomy was assigned based on UNITE v9 dynamic release (Nilsson *et al*., 2019) and Silva 138 (Pruesse *et al*., 2007) databases. The results were then analysed with R (v. 4.0.3), using libraries: phyloseq (McMurdie and Holmes, 2013), tidyverse (Wickham *et al*., 2019), microeco (Liu *et al*., 2021), microbiomeMarker (Cao *et al*., 2022), NetCoMi (Peschel *et al*., 2021). First, for ITS data, we only used ASVs identified to Fungi and Stramenopiles; and for 16S - Bacteria and Archaea. Then, ITS data was repeatedly rarefied at 6,000 and 16S at 11,000 reads using mirlyn package (McMurdie and Holmes, 2014; Cameron *et al*., 2021). We used supervised machine learning model to evaluate the fungal biomarkers of each niche for healthy and infected samples with Supervised Machine Method (SVM), using Total Sum Scaling normalization method in microbiomeMarker R library (Cao *et al*., 2022). We employed Effective Number of Species (ENS) (Jost, 2006) and Simpson index on ASV level to evaluate the dominance of taxa and diversity in each niche. We then used Linear Mixed-Effect Models while controlling for the plant to compare differences between control and infected samples. For the beta-diversity analysis, we used PCoA ordination method based on the weihghthed UniFrac and Bray method for bacteria and fungi. We also used adonis2 (Oksanen *et al*., 2007) and permutest to compare centroid localisation and dispersion between infection groups. To estimate the migration of the bacterial and fungal communities from the niches (sources) to other niches (sinks), we employed SourceTracker (Knights *et al*., 2011). This Bayesian method models the proportion of a microbial community in a given sample (sink) that originates from predefined source environments based on shared microbial taxa. We also used RCbray (Bray-Curtis dissimilarity based Raup-Crick metric) and βNTI (β Nearest Taxon) to evaluate the ecological processes that dominated the bacterial assemblies (Stegen *et al*., 2013; Liu *et al*., 2017), using microeco (Liu *et al*., 2021) R library. βNTI index measures how the differences in species composition between the two samplings differ from what would be expected by the null model of random assembly. When βNTI fell between |2|, microbial assembly was dominated by stochastic mechanisms, and higher than |2|, by deterministic. We then compared the βNTI indexes between control and pathogen-inoculated samples using Wilcoxon signed-rank test. The phylogenetic tree is essential to perform this analysis, we did this only for bacterial communities, as the ITS region is characterised with the variable length for different taxa. Finally, we constructed microbial networks with 200 most frequent taxa and SPIEC-EASI (Sparse Inverse Covariance Estimation for Ecological Association Inference) method (Kurtz *et al*., 2015), using glasso (Graphical Least Absolute Shrinkage and Selection Operator) approach within NetCoMi R library (Peschel *et al*., 2021). The dataset was resampled 30 times and 20 penalty levels with 1e-4 as the smallest ratio, taking the average of symmetric edges. Fast greedy clustering method was used for network visualization.

## Results

### Fungal isolates

Fungal ITS sequences, obtained from symptomatic strawberry leaves (Figure 1B) showed 100% similarity to those of *P. lythri* already present in the GenBank database and phylogenetic tree revealed clustering of the sequences with other representatives of *P. lythri* and *Pilidium concavum* (former name of the *P. lythri* (Marin-Felix *et al*., 2017)) (Figure 1A). The strains collected after 7 days in 23°C in the darkness on Potato Dextrose Agar were flat and granulose, cinnamon in colour (Karimi *et al*., 2016; Marin-Felix *et al*., 2017) (Figure 1E). The presence of hyaline conidia with smooth walls and slightly pointed tips were confirmed under the microscope (Figure 1F). After the inoculation of the surface-sterilized strawberry leaves with the spore suspension and incubation in 24°C in darkness for 7 days, 100% of the leaves showed brown necrotic lesions. Leaves submerged in water did not show symptoms of the disease (Figure 1C). Growth of the cinnamon-colour fungal strains were observed after incubation (24°C for 7 days) of infested leaves on the Petri dishes with the PDA medium, fulfilling he Koch’s postulate (Figure G).

**Figure 1.**
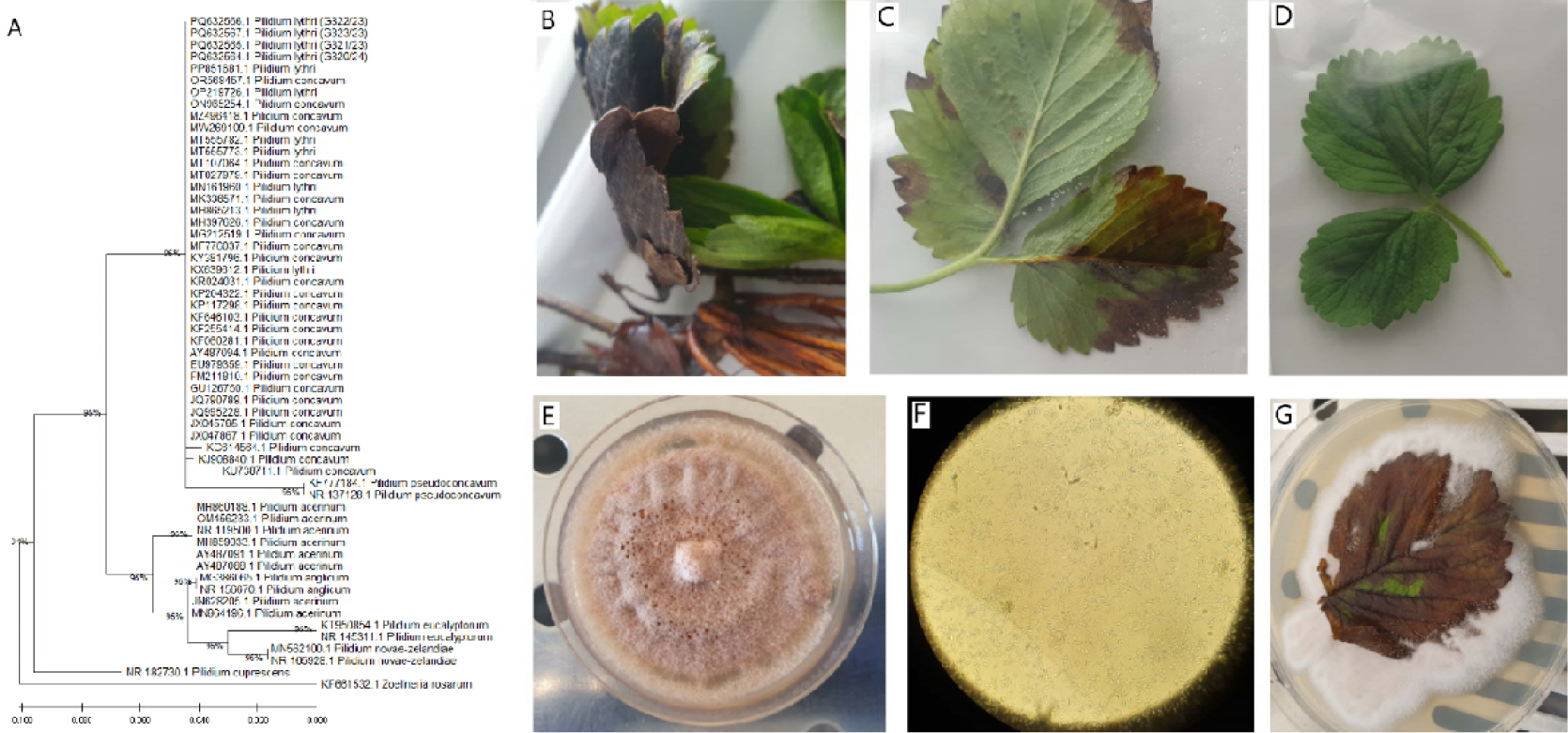
A) Phylogenetic tree of the evolutionary history of *P. lythri* was inferred by using the Maximum Likelihood method and Tamura-Nei model (Tamura & Nei, 1993) and the e tree with the highest log likelihood (−1144.51) is shown. The percentage of trees in which the associated taxa clustered together is shown next to the branches. B,C) Symptoms of the *P. lythri* infection on the strawberry leaves and in the D) control leaf. E) Growth of *P. lythri* after 7 days of growth in 24 °C on PDA medium, F) conidia of *P. lythri* (40x), and G) growth on PDA medium with infected leaf.

### Chemical sensitivity

To evaluate *P. lythri’s* sensitivity to various chemicals, BIOLOG PM 21-25 plates were we incubated with the spore and mycelium solution of three fungal strains (G321/23, G322/23, G323/23) at 24°C and 30°C for 96 hours. An evaluation of the GLLM model revealed that, as expected, the temperature increase was positively correlated with fungal metabolic activity growth, and the substance concentration had negative correlation with the fungal metabolic activity. We observed different response of one of strains - G322/23 that was characterised with higher metabolic activity compared to other 2 (Supplementary Figure 1, Supplementary Table 2). We also evaluated substances that negatively impacted metabolic activity (Figure 2). Top inhibitors included thiourea, dodecyltrimethyl ammonium bromide, tetrazolium violet, alexidine and fluorodeoxyuridine, although the effect size was very small: −0.09, underlining high tolerance of the pathogen to various chemicals.

**Figure 2.**
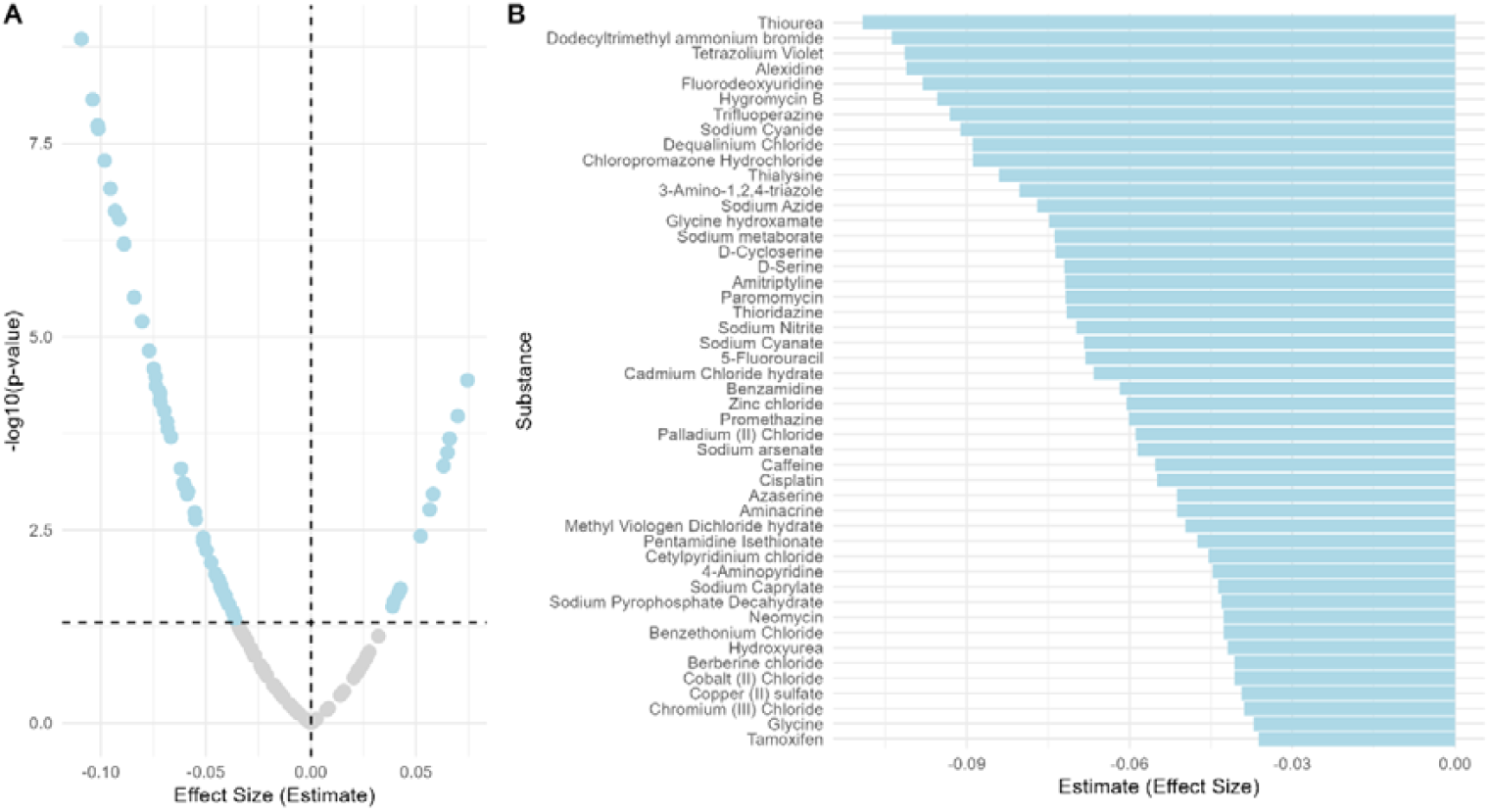
BIOLOG PM21-25 results of the substances that differently influence growth of *P. lythri*. A) Volcano plot visualizing biological and statistical effect for different substances. Gray circles represent no significant importance, light blue - significant results. Dashed vertical lines represent p.value of 0.05 and estimate of -log10(0.05). B) Bar plot representing estimate for each substance significantly inhibiting pathogens metabolic activity in tested conditions.

### Fungal and bacterial community changes after *P. lythri* inoculation of strawberry

Fungal and bacterial molecular markers for 72 samples from soil (24), rhizosphere (24), and roots (24) were amplified and sequenced. Rarefaction curves plateaued for all samples, indicating sufficient sequencing depth. To evaluate the presence of *P. lythri* in infected samples, we employed a predictive approach to identify 15 most important bacterial (Supplementary Figure 3) and fungal (Figure 3) taxa that differentiate healthy and infected strawberry compartments. The analysis revealed that the *Pilidium* fungi belonged to the most important taxa that differentiate between healthy and infected plants and soil (Figure 3). This, along with the visible symptoms of the disease, confirmed that the infection was successful and the pathogen migrated from the soil to rhizosphere and roots.

**Figure 3.**
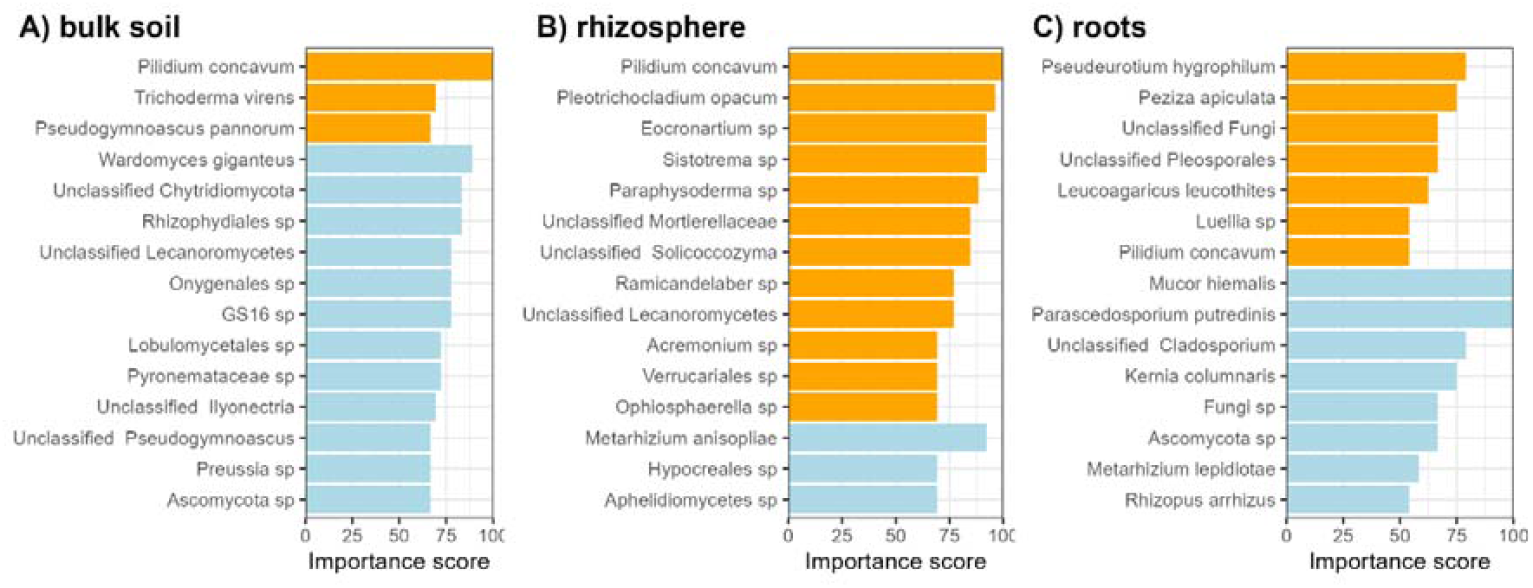
Top 15 fungal taxa found more abundant in (orange) pathogen-inoculated and in (blue) healthy soils and plants identified based on Supervised Vector Machine machine learning approach.

After evaluating that infection was successful, we conducted statistical analysis (LMER modelling while controlling for plant/pot) to compare the alpha diversity of bacterial and fungal communities between healthy and pathogen-inoculated niches. No significant differences in dominance nor richness were observed between healthy and pathogen-inoculated niches at both sampling times (p > 0.05) for bacteria and fungi (Figure 4A, B and Supplementary Table 3). We only noted decreased diversity of fungal microbiome in pathogen-inoculated rhizosphere samples in fungal metacommunities at the second sampling time (p < 0.05). We also conducted beta-diversity analysis of the bacterial (Figure 4 C) and fungal (Figure 4 F) communities. We did not see differences between centroid localization for bacteria (PERMANOVA: r^2^: 0.019, p > 0.05), but the groups were characterized with different dispersions (permutest: F = 0.0011, p < 0.05), suggesting increased variability in response of bacterial community to pathogen infection. In fungal metacommunities, the analysis revealed no significant differences between infected group and the control (PERMANOVA: R^2^ = 1.29, p > 0.05, permutest: F = 0.77, p > 0.05) (Supplementary Table 3).

**Figure 4.**
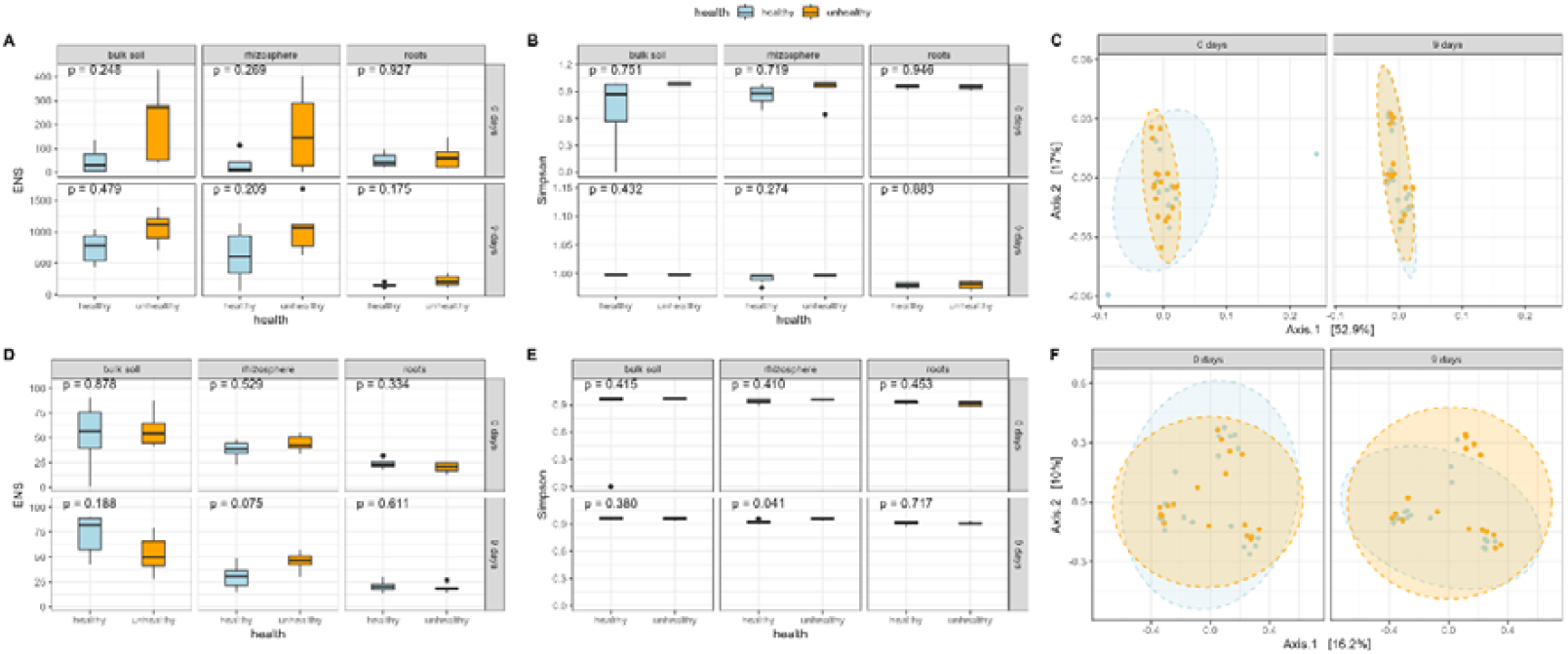
Comparison of alpha and beta diversity of bacterial (A, B, C) and fungal (D, E, F) communities of strawberry niches immediately after *P. lythri* inoculation (0 days) and after 9 days of (9 days). LMER model was used for comparison of non-independent samples in alpha diversity. Light blue color represents samples without pathogen inoculation, and orange - with inoculation.

To elucidate the influence of *P. lythri* on microbial migration between the niches, we used samples from sampling at 0 days as a source, and samples from sampling at 9 days as sinks. In bulk soil, only bacterial community revealed significantly higher share of microorganisms in bulk soil and a lower portion from unknown source in infected samples (Figure 5A). Furthermore, we observed migration of bacteria from bulk soil to rhizosphere for both bacteria and fungi in infected communities, but also from rhizosphere to rhizosphere for bacteria in infected communities. Finally, analysis revealed increased migration from rhizosphere to roots in infected samples in bacterial community.

**Figure 5.**
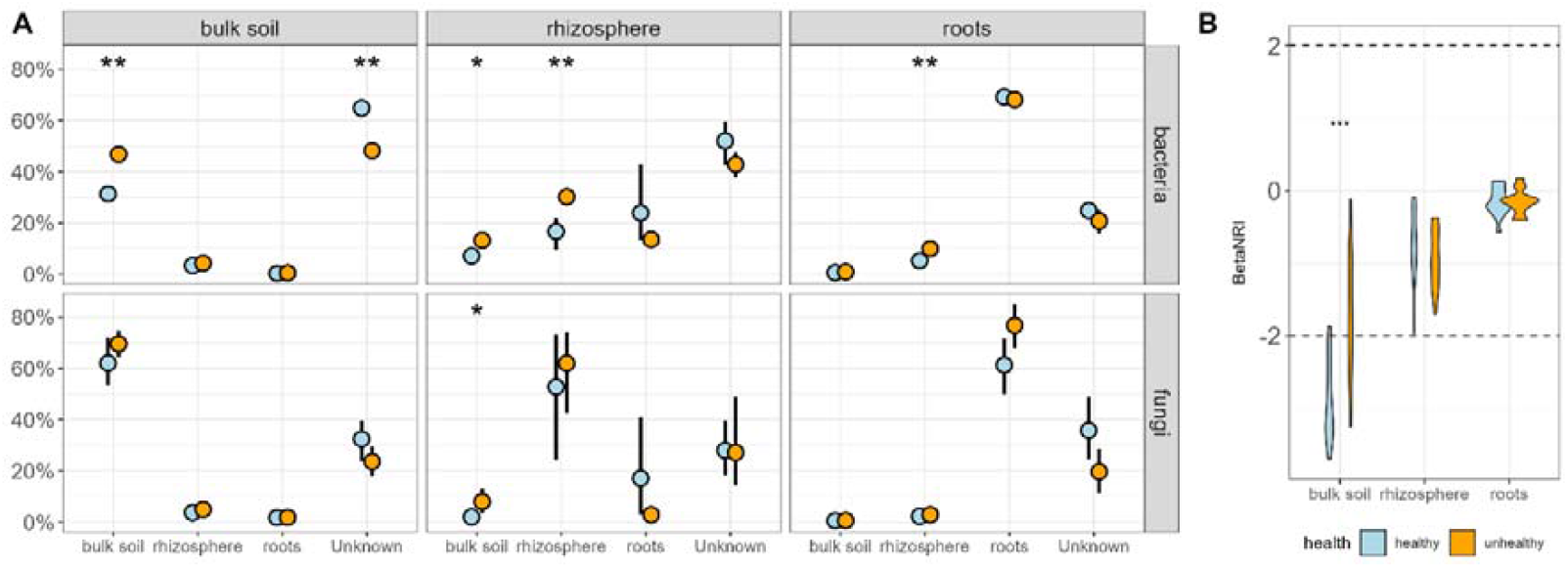
Source tracking analysis (A) of bacterial and fungal microbiomes between sampling one (source) and two (sink). Comparison between healthy (light blue, left) and contaminated (light orange, right) samples. Values are shown as mean standard error from bootstrap. (B) BetaNRI index analysis of bacterial community in second sampling. Significant differences on both panels between healthy and contaminated samples are shown with asterisk.

We also evaluated the mechanisms involved in bacterial community assembly by calculating the betaNRI index for bulk soil, rhizosphere, and root samples collected during the second sampling period (Figure 5B). The results revealed a dominance of stochastic processes in both healthy and infected rhizosphere and root samples of strawberry (betaNRI values between −2 and 2). Furthermore, we observed differences in community assembly processes between healthy and infected bulk soil, with a shift from deterministic processes (characterized by lower turnover) in healthy bulk soil to a dominance of stochastic processes in infected soil.

Finally, to evaluate the influence of *P. lythri* on the structure of fungal community, we compared the network structure in the rhizosphere niche between control and infected samples during the second sampling (Figure 6, Supplementary Figure 3, Supplementary Table 5). Both networks showed similar characteristics in terms of the number of components (1), clustering coefficient (0.995, 0.999), modularity (−0.007, −0.008), edge density (0.994, 0.999) and natural connectivity (0.375, 0.310), average dissimilarity and average path length (0.6, 0.6). On the other hand, the control rhizosphere exhibited a higher percentage of positive edges (52.308% vs. 43.344%) but lower vertex and edge connectivity (58, 71).

**Figure 6.**
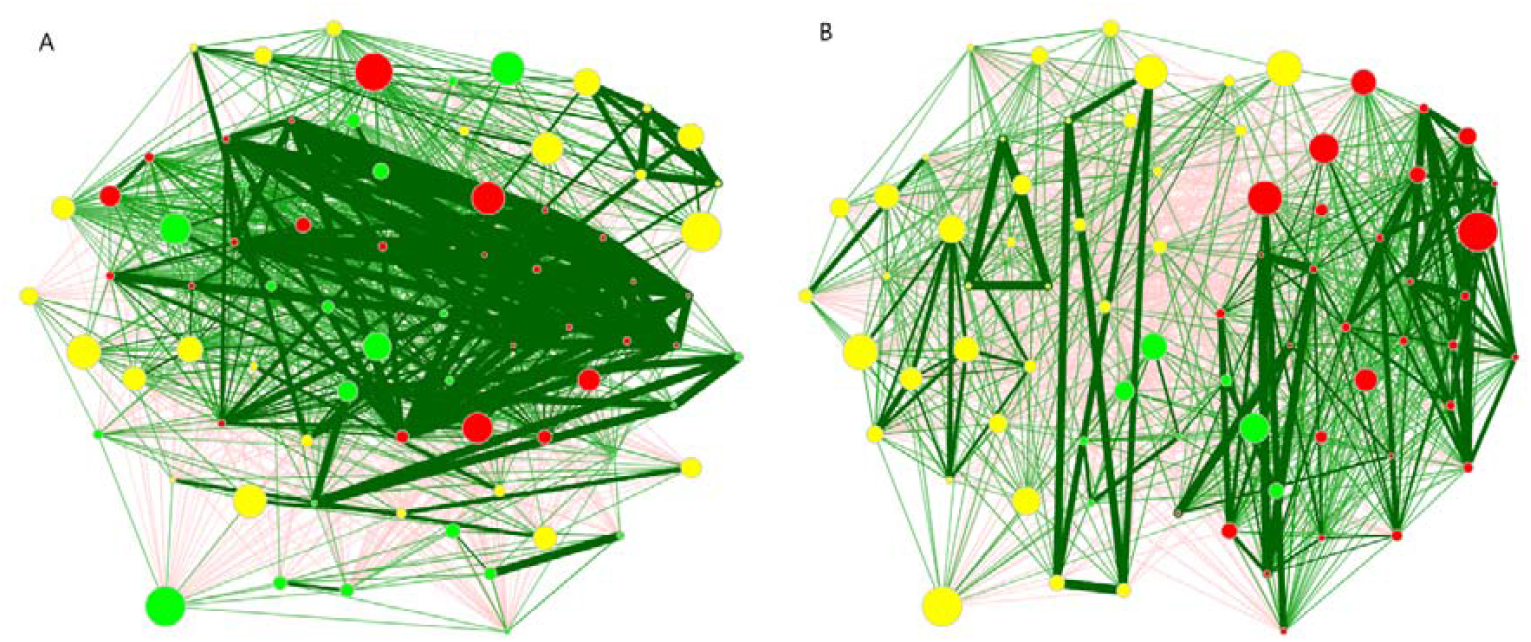
Fungal network analysis in A) control and B) pathogen-inoculated rhizosphere samples of strawberry. Each node represents individual fungal ASV, and node size is proportional to number of reads. Node colours represent the same clusters. Green lines represent positive interactions and red-negative. The position of the same nodes is the same in both networks.

Notably, changes were observed in centrality of networks (Jaccard Index). Degree, betweenness and hub taxa measures showed no overlap in most connected, keystone and key hub taxa, significant shift was observed in closeness of most central nodes (Table 1).

**Table 1.** Centrality changes between networks of rhizosphere samples in control and *P.* contaminated samples. Asterisk shows statistically significant change.

|  | Degree | Betweenness | Closeness | Eigenvector | Hub taxa |
| --- | --- | --- | --- | --- | --- |
| Jaccard Index | 0 | 0 | 0.071 | 0.209 | 0 |
| p-value (P≤Jacc) | 1 | 0.296 | 0.027* | 0.055 | 1 |

Importantly, *Pilidium* was identified as a connector taxa in the control rhizosphere (betweenness = 1), loosing this status in pathogen-inoculated (betweenness = 0) samples (Supplementary Table 5).

## Discussion

Here, we investigate the underexplored role of emerging thermotolerant fungal pathogen in shaping plant microbiomes using newly isolated strains of *Pilidium lythri*. We assessed its thermal tolerance by screening metabolic response to 120 different chemical substances and explored its impact on the structure, assembly processes and association networks of bacterial and fungal metacommunities across strawberry soil and plant niches (bulk soil, rhizosphere, roots).

*In vitro* evaluation of the impact of various chemical substances on the growth and development of the tested phytopathogen at two temperatures (24 and 30 °C) using BIOLOG microarrays raised serious concerns, as *P. lythri* demonstrated overall higher metabolic activity at higher temperatures. This aligns with the growing evidence of global warming contributing to the spread of fungal phytopathogens to new areas (Ward *et al*., 2007; Bebber *et al*., 2013; Chaloner *et al*., 2021). Among the 120 chemical substances tested, top five presented inhibition mechanism on pathogens metabolic activity, likely by disruption of electron chain, destabilization of cell membranes, and inhibition of DNA/RNA synthesis (Waldorf and Polak, 1983; Mamouei *et al*., 2018; Ansari *et al*., 2024; Yang *et al*., 2025). Unfortunately, although significant, the effect of the inhibition was modest. These results underline high resilience of the pathogen that points to its high environmental tolerance and potential to spread.

While the dissemination of fungal pathogens is widely reported (Bebber *et al*., 2013), still little is known about the effect of such invasive microorganisms on the microbial structure of the plant holobiont. A number of studies have reported significant differences in the α-diversity of microbiome structure between healthy and infected soil and plant niches (Lefcheck *et al*., 2015; Delgado-Baquerizo *et al*., 2016; Bakker *et al*., 2018). In the present study, we compared alpha and beta diversity of bacterial and fungal metacommunities in healthy and *P. lythri*-infected bulk soil, rhizosphere, and roots at two time points: directly after pathogen inoculation and 9 days post-inoculation. For the bacteria, alpha diversity did not differ between inoculated and control samples, but we observed increased community dispersion, that points to possible destabilization of bacterial assemblages under biotic stress. To test this, we analyzed the mechanisms involved in bacterial microbiome assembly. The results indicated that composition of bacterial communities in healthy and diseased rhizosphere and roots samples was shaped by stochastic processes (i.e. random colonization, dispersal, and ecological drift). Interestingly, we observed a shift from deterministic processes in healthy bulk soil, to stochastic processes in *P. lythri*-infected samples, indicating a disruption in microbial turnover. Infected soil showed random processes that could have been triggered by disturbances in microbial community caused by the infection, described by the Anna Karenina Principle (AKP) (Arnault *et al*., 2023). Analysis of microbial migration suggests increased migration of bacteria from the soil in pathogen-inoculated samples, as well as increased migration of bacteria from the rhizosphere to the roots in infected plant samples. This inconsistency between migration (suggesting plant filtering) and microbial assembly (suggesting random processes) could be caused by the smaller scale of deterministic processes being masked by larger-scale microbial community assembly. Furthermore, these two mechanisms represent different time scales, where βNRI focuses on broader, longer-term assembly processes (Arnault *et al*., 2023). Importantly, migration of microorganisms can be caused by random dispersal (stochastic processes) but may appear deterministic as migration increases. Thus, these two processes are not necessarily contradictory.

In the fungal microbiome, we observed increased migration of fungi from bulk soil to the rhizosphere in the inoculated samples compared to control, highlighting the importance of this niche in fungal microbiome dynamics and the role of plant in microbial recruitment under pathogen stress (Bais *et al*., 2006; Berendsen *et al*., 2012). The alpha diversity results revealed an increased diversity of the fungal community under pathogen influence in the strawberry rhizosphere, and we then compared the microbial networks in this niche to further explore the influence of the pathogen. A number of methods for construction and analysis of co-occurrence networks had been developed, and SPIEC-EASI is characterized with important advantages, such as accounting for compositional type of data and mitigation of non-independence of samples. Also, an implementation of glasso approach allowed for the analysis in high-dimensional setting with more variables than samples (Kurtz *et al*., 2015). Thus, we implemented these methods for the analysis in current study. Comparison of healthy and infected samples revealed shift in microbial fungal networks under *P. lythri* influence. The fungal networks in infected rhizospheres exhibited lower percentages of positive edges compared to control, and lower robustness of networks, suggesting restructuring of fungal community in response to pathogen presence. These findings underline decrease in natural connectivity percentage of positive edges, that suggest lowered positive interactions between fungal taxa. Infection with the pathogen significantly disrupted fungal networks, by decreasing networks connectivity, which indicates a shift toward increased competition and antagonism within the fungal microbiome. This suggests adaptation of the fungal meta-community, making it more resistant to further infections and environmental stressors (Hernandez *et al*., 2021).

Concluding, in this study, we isolated and molecularly identified isolates of spreading fungal phytopathogen from symptomatic strawberry plants - *P. lythri*. We observed pathogen’s high tolerance to various chemical substances in both, lower and higher temperature, highlighting it hight thermal plasticity and resilience. We also noted significant variability in tolerance caused by chemical substances between strains. We used *P. lythri* to evaluate ecological impacts of spreading fungal phytopathogen on microorganism metacommunities in bulk soil, rhizosphere and roots of strawberry with high-throughput marker sequencing. We revealed that *P. lythri* infection had a major effect on both the structure and dynamics of the plant associated microbiome. Its ability to shift bacterial microbial assembly mechanism from deterministic to stochastic in the bulk soil, underscores its capacity to destabilize plant-associated microbiomes. Additionally, we observed that *P. lythri* shifted the fungal microbiome dynamics in the strawberry rhizosphere towards reducing mutualistic connections, while maintaining overall network stability. This underlined adaptive reorganization of fungal community, potentially increasing resilience to secondary infections and other (a)biotic stresses.

These findings underline the critical role of emerging, thermotolerant fungal pathogens like *Pilidium lythri* in destabilizing soil microbiomes and plant-microbiome interactions. This is significantly important in the context of global warming, which accelerates spread and impact of such pathogens.

## Author Contribution

Conceptualization: DS, MF, JP, Data curation: DS, JP, Formal analysis: DS, Funding acquisition: DS, Investigation: DS, JP, Methodology: DS, JP, MF, EH, Project administration: DS, Resources: MF, DS, EH, Software: DS, JP, EH, Supervision: DS, Validation: DS, Visualization: DS, Writing – original draft: DS, Writing – review & editing: DS, EH, JP, MF

## Acknowledgments

This research was funded in whole by National Science Centre, Poland, contract number: 2022/45/N/NZ9/02089. For the purpose of Open Access, the author has applied a CC-BY public copyright license to any Author Accepted Manuscript (AAM) version arising from this submission. The DS’s 3-month visit at the Institute of Environmental Sciences, Leiden University, Leiden, Netherlands was financed by the Polish National Agency for Academic Exchange, contract number: BPN/BEK/2023/1/00125/U/00001.

## Data Availability Statement

Metabolic profile data and R code used for ststistical analysis were deposited at gihub (https://github.com/dsiegieda/biolog_and_pot_Pilidium) for peer review purposes. After acceptance of the manuscript for publication, this data will be published in RepOD repository (https://repod.icm.edu.pl/), under permanent doi number. ITS sequences of 4 *Pilidium lythri* strains were deposited in GenBank under accession numbers: PQ632564.1, PQ632565.1, PQ632566.1, PQ632567.1. Metataxonomic sequencing data of soil and plant niches were deposited at SRA database under accession numbers: PRJNA1213298 for 16S and PRJNA1213307 for ITS. Benefits generated: benefits from this research accrue from the sharing of our data and results on public databases, as described above.

## Declaration of generative AI and AI-assisted technologies in the writing process

During the preparation of original draft of this manuscript the author used ChatGPT in order to improve language and readability. After using this tool, the author reviewed and edited the content as needed and take full responsibility for the content of the publication.

